# Dissecting mammalian cortical circuit development at single-cell resolution using inducible barcoded rabies virus

**DOI:** 10.1101/2025.11.07.687321

**Authors:** Zijian Zhang, Brooke R. D’Arcy, Lewei He, Diego Arroyo, Samantha Deasy, Elizabeth A. Matthews, Zihan Yan, Nirvika Rastogi, Walter Mancia Leon, Kiran Shehnaz Kaur, Derek G Southwell, Z Josh Huang, Debra L. Silver, Dmitry Velmeshev

## Abstract

Highly organized circuits of connected neurons enable diverse brain functions. Improper development of these circuits is associated with neurodevelopmental disorders, and understanding how circuits are formed is crucial for unraveling the mechanisms of these diseases. We currently have an incomplete picture of how specific brain circuits develop and how they are affected in disease, because we lack methods to study them at scale and with single-cell resolution. Monosynaptic rabies tracing is the gold standard method to study circuit architecture. However, it suffers from cellular toxicity, low throughput, lack of control over the timing of labeling, and the inability to access the molecular profiles of individual neurons. To address these issues, we developed an inducible barcoded rabies virus (ibRV) to enable temporal-controlled labeling of synaptic circuits followed by high-throughput single-cell genomics readout. ibRV allows for dissecting neuronal circuit changes over time at single-cell and spatial resolution. We applied ibRV to study the development of specific mouse cortical circuits during late prenatal and postnatal life using single-cell genomics and unbiased spatial transcriptomics as readouts. We characterized and quantified developmental connectivity patterns and molecular cascades that underlie their formation. Additionally, we constructed functional *in silico* circuit models that enable interrogation of circuit function and dysfunction at specific developmental stages. Our study provides novel tools for circuit analysis and can provide new insights into the mechanisms of mammalian brain development.

## Introduction

During late embryonic and early postnatal development, neural circuits undergo a rapid transformation from broadly permissive connectivity into precisely tuned, functional networks^1^. Axonal projections complete their final pathfinding and target selection, followed by activity-dependent refinement that eliminates exuberant or inappropriate contacts while stabilizing behaviorally relevant ones^2,3^. Distinct circuit modules mature along unique trajectories, for example, thalamocortical sensory pathways refine earlier than intracortical associative networks^4^, and excitatory synaptogenesis precedes the maturation of parvalbumin-mediated inhibition required for precise spike timing^5^. Each of these developmental programs is tightly scheduled; perturbing maturation or impaired sensory pathway refinement has been causally linked to neurodevelopmental disorders such as autism spectrum disorder^6,7^, intellectual disability^8^, and epilepsy^9,10^. How specific cell types wire into evolving networks, which neurons connect, and when connections form, remains a central challenge in developmental neurobiology^11^.

Monosynaptic rabies virus (RV) tracing has become a cornerstone methodology for resolving synaptic connectivity with genetic and single-synapse specificity^12–15^. The emergence of barcoded rabies, coupled with single-cell sequencing, has further transformed classical circuit tracing into a high-throughput connectomic platform, enabling quantitative profiling of synaptic inputs across the brain^16–20^. However, the utility of RV in developmental and longitudinal studies remains fundamentally constrained by its cytotoxicity^21–24^: glycoprotein-complemented starter neurons support persistent viral transcription and replication, driving rapid disruption of intrinsic membrane properties, reduced firing stability, innate immune activation, and transcriptional reprogramming^25–29^. These effects culminate in widespread neuronal death within 7–14 days post-infection, constraining the use of conventional rabies virus systems for studying developmental events or dynamic circuit changes.

To overcome these limitations, we developed an inducible barcoded rabies virus system that retains high-throughput molecular readout while markedly reducing neuronal toxicity and enabling precise temporal control of viral replication. We engineered destabilizing domains (DD^30^ and DHFR^31^) into the rabies phosphoprotein (P) and/or polymerase (L), providing tunable, small-molecule–regulated control over rabies transcription. This strategy minimizes viral burden during non-induced periods yet preserves robust barcode expression compatible with single-cell genomics.

By combining in utero cortical ventricular delivery with temporally programmable induction, this platform enables circuit labeling across the entire brain during late embryonic and early postnatal development. When coupled with single-nucleus RNA-seq and spatial *in situ* single-cell sequencing, it provides a scalable framework to profile the dynamic maturation and remodeling of synaptic networks, linking structural rewiring to molecular state transitions in defined neuronal subtypes. Together, ibRV achieves low toxicity, temporal control, and genomics-level scalability, empowering systematic dissection of brain-wide circuit assembly during critical developmental periods.

## Results

### Construction of an inducible rabies virus system

Rabies virus toxicity arises from three main factors: toxic viral proteins, immune activation, and depletion of host cell resources^32–34^. We reasoned that by controlling the expression of proteins that play key roles in virus replication and transcription (N, P, or L), we can reduce viral protein levels, thereby lowering toxicity, immune responses, and metabolic burden. In order to generate an inducible rabies virus system, we tested two small-molecule–regulated protein destabilization systems for inducible control of target protein levels. One is based on a mutated FKBP12 domain stabilized by the compound Shield1^30^, and the other on *E. coli* DHFR stabilized by trimethoprim (TMP^31^). Both small molecules cross the blood–brain barrier, enabling *in vivo* regulation of protein expression^35–38^. We then fused a molecular switch to either their N- or C-terminus of N, P, and L, and constructed pairwise combinations, yielding 24 constructs (**Fig. s1A&B**). We then tested all 24 constructs in the B7GG cell line used for rabies virus recovery from a plasmid to assess their ability to recover functional viral particles and to determine whether replication could be effectively regulated by the corresponding small molecules (**Fig. 1B&s1C, D)**. The construct with the viral phosphoprotein P fused with DHFR at the N terminus (Pn-DHFR) **(Fig. 1A)** and P&L both fused with DD at N terminus (PLn-DD) demonstrated the best specificity and dynamics of response to TMP treatment (**Fig. 1B**) or Shield-1 separately **(Fig. S1E)**. TMP is an antibiotic with good central nervous system penetration and has been used to treat certain bacterial and parasitic encephalitides, such as toxoplasmic encephalitis^39–41^. Given its extensive study, and ready availability of TMP in both humans and animals, we selected Pn and its paired TMP as our primary candidates, hereafter referring to the Pn-DHFR RV as the inducible rabies virus (iRV). Fluorescence of mCherry encoded by iRV quickly disappeared after iRV infection without TMP, but in the presence of TMP virus replicated and infected neighboring cells. Additionally, we are able to perform viral reactivation by decoupling virus infection and TMP activation by as long as 10 days (**Fig. 1C &s1E**). Importantly, we did not observe Pn-DHFR virus spread in the absence of TMP, even after continuous passaging of infected cell lines for three months (**Fig. s1F**). It is likely due to the fact that any protein-truncating mutation in DHFR at the N terminus disrupts expression of the essential viral protein P. This feature of Pn-DHFR RV is aligned with SiR-N2c self-inactivating rabies^28^, which prevents the potential loss of the inactivation domain during viral amplification^42,43^. Therefore, iRV enables small-molecule-switch–mediated control of viral replication.

**Figure 1.**
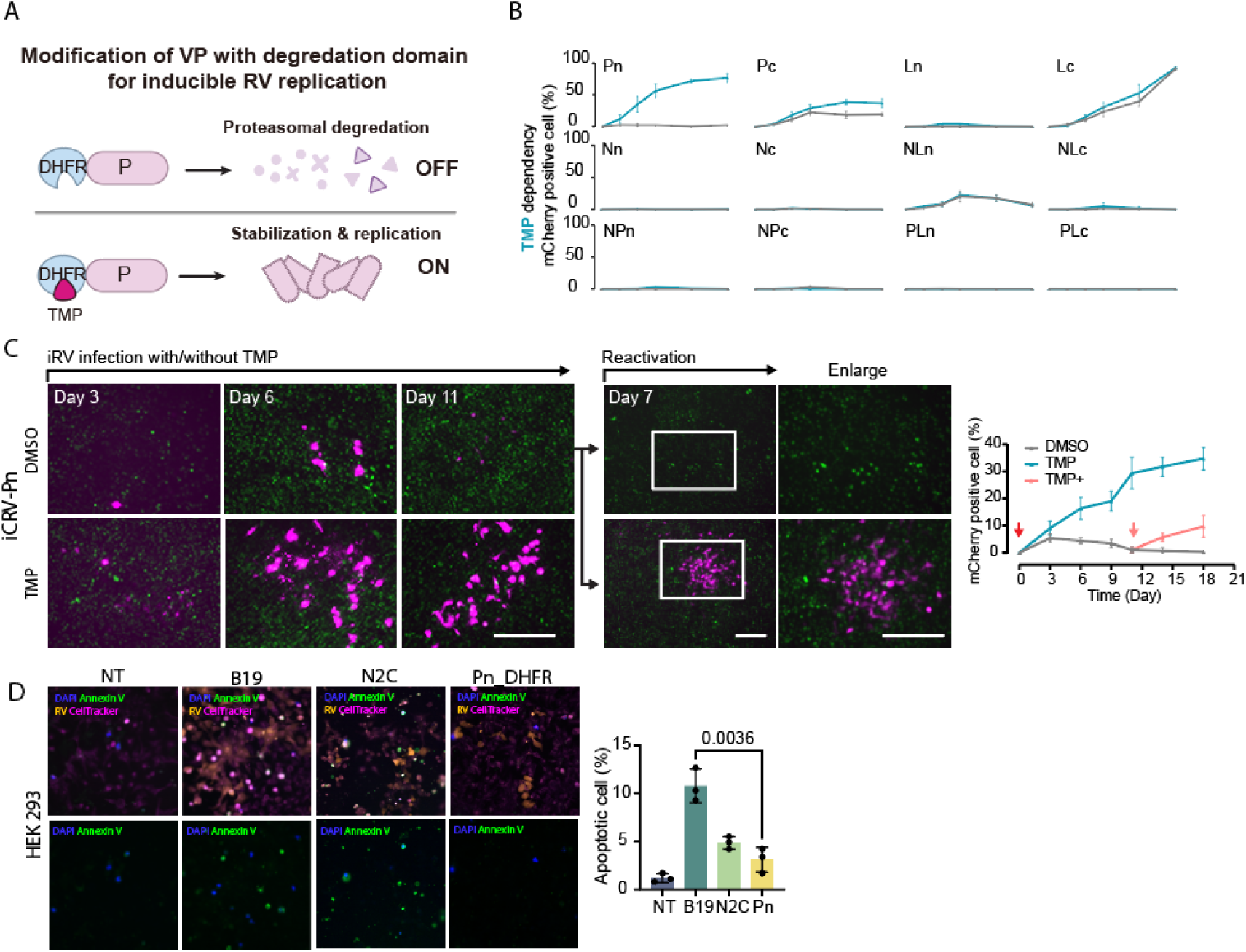
iRV through fusion with a DHFR achieves inducible proliferation and toxicity reuction. A. Fusion of viral proteins with the DHFR molecular switch enables control over their degradation and stabilization. B. Screening of 12 DHFR–viral protein fusion design strategies. C. iCRV-Pn-DHFR controlled proliferation and reactivation 10 days post initial infection. D. Cytotoxicity test with Annexin VI assay *in vitro*.

### Validation of toxicity and neuronal circuit labeling of iRV *in vitro*

In order to test whether control of protein P level leads to reduced cell toxicity, we generated viral stocks of iRV, the first-generation B19 rabies, as well as less toxic CVS-N2c strain of rabies^27^. We used these versions of monosynaptic RV to infect non-neuronal HEK293, as well as neuronal mouse Neuro2a and human neuroblastoma SHSY-5Y cells. We then utilized immunostaining (IHC) and fluorescence-activated cell sorting (FACS) to quantify the number of apoptotic cells using Annexin V & DAPI assay. iRV demonstrated 3-fold lower cell death than B19 RV and toxicity comparable to N2c RV (**Fig. 1D&s2A**) *in vitro*, at 4 days post infection. Additionally, we demonstrated the ability of iRV to spread throughout neuronal circuits in a TMP-dependent manner *in vitro in primary cultured hippocampal neurons* and *ex vivo* using organotypic mouse brain slice culture (**Fig. s2B&C**). Therefore, iRV demonstrates lower toxicity than B19 RV and a level of toxicity reduction comparable to N2c RV *in vitro*, while preserving transsynaptic transmission.

### Reduced cytotoxicity of iRV in labeling hippocampal neurons in mice

To test the ability of iRV to label neuronal circuits *in vivo*, we utilized a widely adopted rabies tracing transgenic mouse line RphiGT that expresses TVA receptor and rabies glycoprotein G in a Cre-dependent manner. This mouse line was crossed to the Cre-dependent ZsGreen line to enable fluorescent readout of Cre activity (**Fig 2A**). We produced iRV, B19 and N2c rabies encoding Cre and injected them into the hippocampus CA3 region (6 weeks old) mice. The brain was analyzed 1 or 2 weeks after rabies injection using IHC for ZsGreen and RV-encoded mCherry (**Fig. 2B**). Brain of animals in the B19 group contained few red or green cells, consistent with high toxicity of this rabies strain that is expected to kill most infected neurons after 2 weeks of infection^26^. N2c group displayed several double-positive cells. In the iRV group at 2 weeks, the majority of cells were positive for ZsGreen but not mCherry, indicating that iRV induced target locus recombination and mostly cleared out from the cells. Compared with the 1-week time point, the number of surviving iRV-labeled neurons at 2 weeks was comparable, and remained significantly higher than that in the B19 group, at a level similar to N2c (**Fig. 2C**). Therefore, iRV enables inducible and low-toxic labeling of neuronal circuits, such as during behavioral paradigms, or to transiently deliver gene cargo to synaptically coupled neurons.

**Figure 2.**
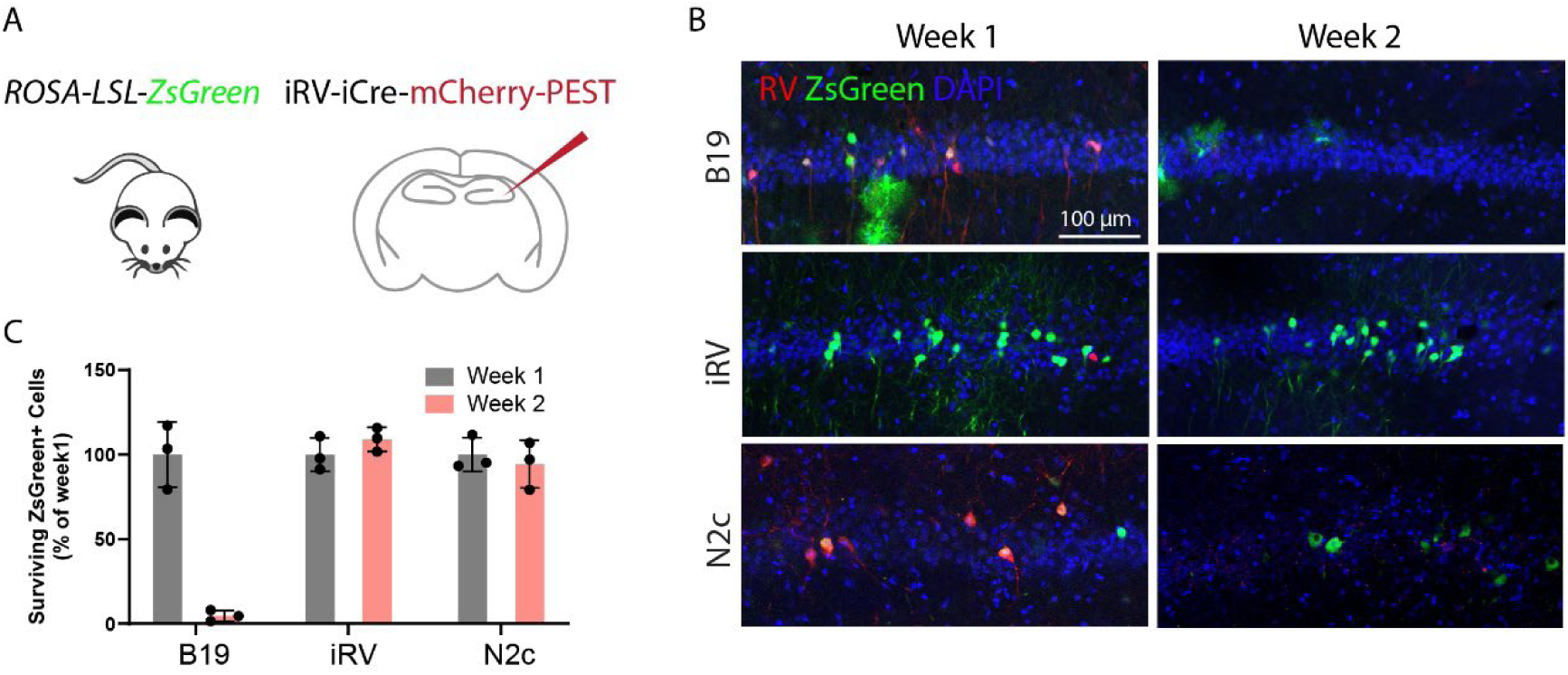
Reduced cytotoxicity *in vivo*. A. Quantification of survival rabies virus infected hippocampal neurons of Rosa-LoxP-STOP-LoxP-ZsGreen mice at 1 week, 2 weeks. B&C. IF showing EGFP⁺ infected cells and mCherry⁺ rabies expressing reporter at 1- and 2-weeks post-injection.

### Large-scale labeling of developing neuronal networks with in utero iRV infection

Above, we demonstrated the ability of iRV to label neuronal circuits in the adult brain. However, injections of RV in the adult brain only label a limited number of neurons in the vicinity of the injection site and do not allow for study of circuit development. To increase the scale of labeled networks and gain access to critical stages of circuit development, we first attempted to inject iRV into the lateral ventricle of neonatal (P0) mice. This approach is based on published data demonstrating whole-brain infection of adeno-associated virus (AAV) when delivered via this route and at this age^44^. To test this hypothesis, we performed P0 injection of RV in the ventricle of mice expressing rabies glycoprotein (G) and TVA receptor for EnvA-pseudotyped rabies in Foxp2-positive striatal and cortical layer 6 excitatory neurons (**Fig. 3A**). However, we observed that RV only labeled a few cells compared to previously published studies using P0 AAV injections^44^. We reasoned that a much larger size of the RV particle (180×75nm)^45^ compared to AAV (20-25nm in diameter)^46^ precludes its diffusion across the brain even at P0. To circumvent this obstacle, we performed *in utero* injection of iRV into cortical ventricles at embryonic day 16.5 (E16.5) when the majority of Foxp2 neurons had been born^47^ (**Fig. 3B**). Since TMP has been reported to cross the placental barrier^48,49^, we treated the pregnant and lactating dams with TMP in drinking water and observed widespread labeling of neurons with iRV in the cortex, striatum and thalamus of pups at P3 (**Fig. 3C**). Without TMP induction we did not observe labeling in the hippocampus or upper cortical layers, consistent with specific infection of Foxp2 starter neurons and the known anatomical locations^47,50^. We induced circuit labeling with TMP at developmental stages between E16.5 and P0, P0 and P7, as well as P7 and P14 (**Fig. 3C**). Importantly, we observed much smaller number of cells labeled at P3 and at P7 and P14, suggesting that the virus does not replicate or spread in the absence of TMP and clears out from the cells withing 10 days. Labeling at P7-P14 was less efficient, possibly many cells losing iRV prior to TMP reactivation at P7 (**Fig. 3C**). Compared to the B19 RV, iRV did not induce astrocyte and microglia activation or apoptosis at P3 as measured by Gfap, Iba1 and CC3 IHC (**Fig. s3**). Therefore, iRV can label circuits at specific developmental time points and at a near whole-brain scale.

**Figure 3.**
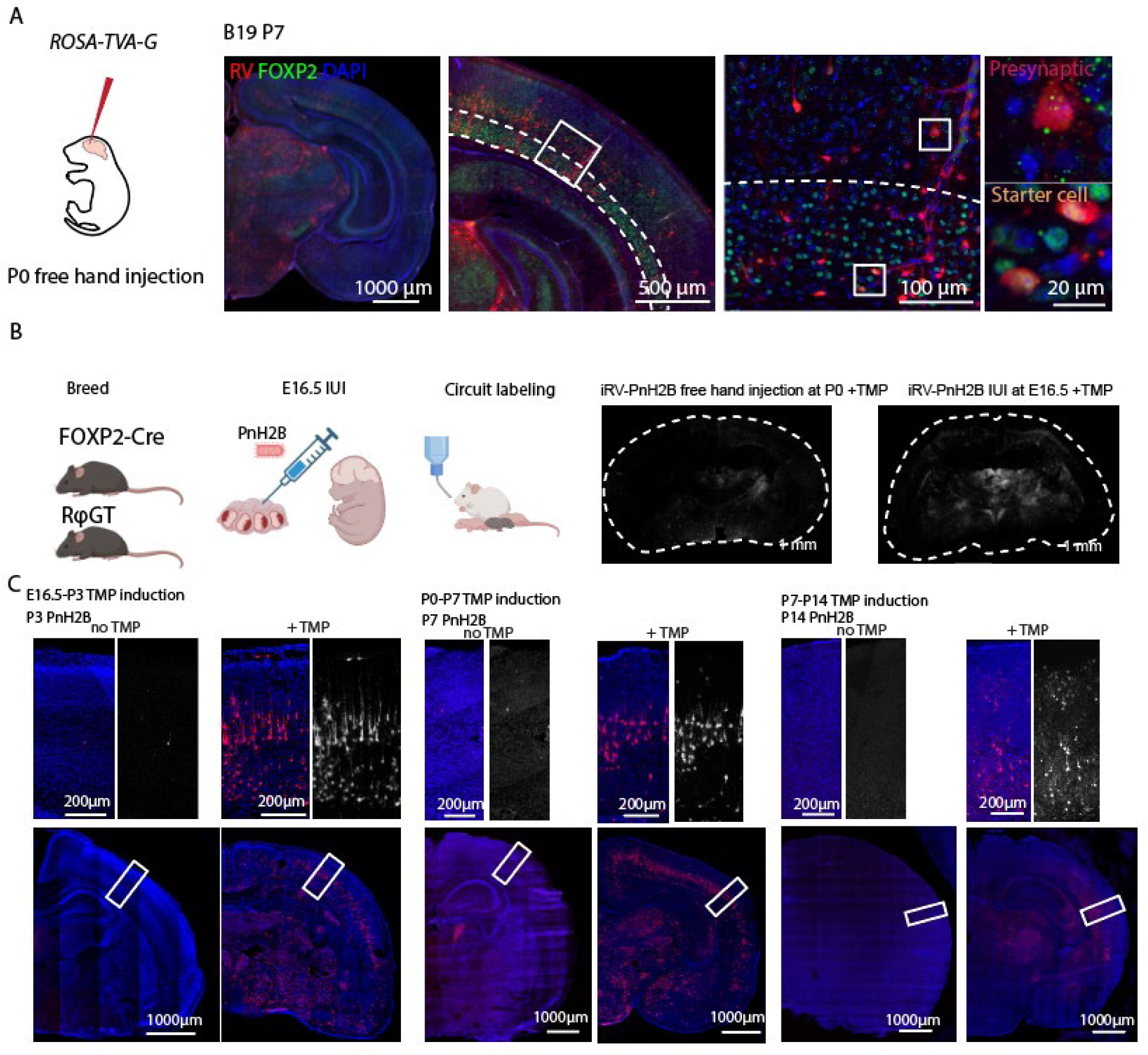
*In utero* injection of iRV results in broad neuronal labeling and flexible initiation at distinct developmental stages. A. Ventricular injection of iRV at P0 labels receptor-expressing neurons in cortical layer VI (Foxp2-Cre x RphiGT). B. In utero injection at E16.5 results in higher labeling efficiency. C. RNAscope detects iRV rabies RNA expression at different developmental stages.

### Production of high-complexity libraries of inducible barcoded rabies virus

Next, we aimed to combine our iRV system with single-cell genomics and spatial transcriptomics to dissect development of specific circuits at single-cell resolution. We have created a high-complexity barcoded rabies virus library based on our inducible system. We cloned a two-segment random barcode sequence following the mCherry gene of our Pn-RV vector, which allows for a hypothetical 10^12^ unique barcodes^18^ (**Fig. 4A&s4A**). We then used NEB 10-beta electrocompetent *E. coli* to amplify the plasmid libraries. Transformed bacteria were plated onto twenty 15-cm LB agar plates, and a small fraction of the transformation mix was serially diluted to estimate colony number, yielding >2 × 10⁶ colonies per preparation (**Fig. 4A**). The resulting amplified plasmid library was used to generate inducible barcoded RV (ibRV) using a cell line we created to improve rabies virus recovery efficiency. Specifically, we used HEK293T cells stably expressing T7 RNA polymerase and rabies glycoprotein via lentiviral transduction, followed by single-cell cloning. Clones were screened for ibRV recovery efficiency in the presence of TMP (**Fig. s4B**). We found that the addition of MG-132 further enhanced the recovery rate of iRV, likely by inhibiting proteasome activity and preventing degradation of viral proteins fused to the destabilizing DHFR domain (**Fig. s4c**). After EnvA pseudotyping of the ibRV library, we isolated viral RNA from and performed deep sequencing of both the plasmid library and the viral library. We observed ∼1.3 million barcodes in the plasmid library and ∼550,000 barcodes in the viral library, suggesting that we can recover more than 40% of the original barcodes during ibRV preparation and that for the majority of barcodes, each barcode is represented by a small number of transcripts (corresponding to UMIs, unique molecular identifiers) (**Fig. 4B**). Consistent with a recent work from Shin et al^20^, recovering iRV in 96-well plates did not effectively increase barcode diversity. These results represent a significant increase in both the number of viral barcodes and their recovery rate compared to a recently published study, where 20-30% of the plasmid barcodes could be recovered in the pseudotyped virus^18^. It is possible that the relatively slow recovery rate of iCRV limited the formation of large super-colonies, thereby indirectly increasing barcode diversity. We have confirmed that iRV can be induced with TMP *in vitro* and *in vivo*. As a result, we were able to combine our novel inducible rabies virus with molecular barcoding to enable dissecting developmental circuit dynamics at single-cell resolution.

**Figure 4.**
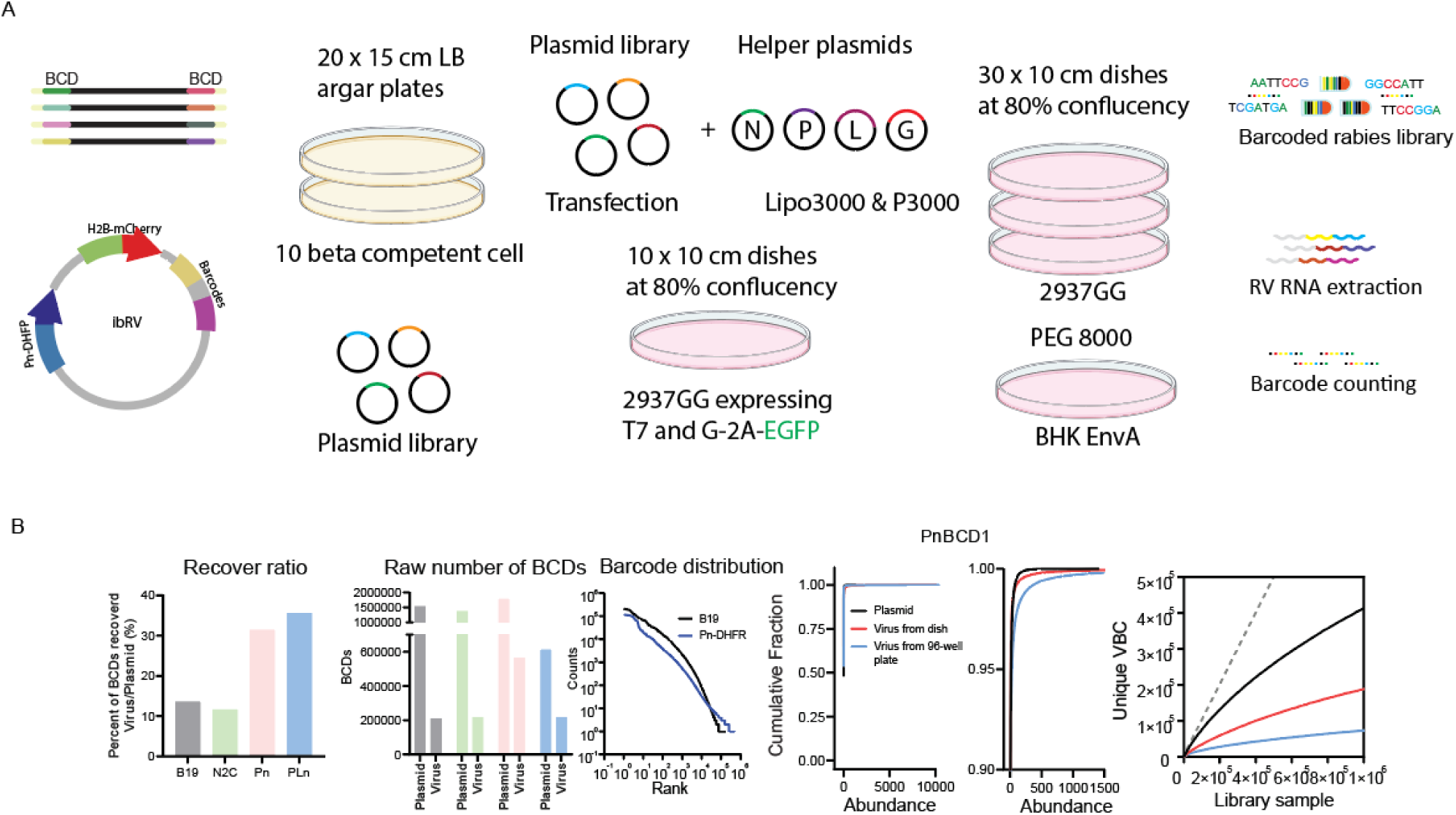
The optimized production approach enables generation of high-diversity barcoded iRV. (A) Schematic illustrating ibRV production by recovering virus in pre-screened 293T-7GG cells, along with barcode diversity quality control. (B) Barcode diversity analysis comparing barcode abundance and distribution across different viral library lines.

### Construction of spatially resolved maps of developing cortical circuits using ibRV and unbiased spatial transcriptomics

Since snRNA-seq requires tissue dissociation to isolate nuclei, it leads to the loss of spatial information of individual neurons within circuits. Spatial organization plays an important role in circuit architecture, and molecular information alone is not always sufficient to resolve neuronal subtypes, especially for subdivisions of subcortical regions such as the thalamus. To resolve cortical circuit development with single-cell and spatial resolution, we combined our ibRV approach with spatial transcriptomics. In order to dissect the developmental formation and refinement of thalamocortical and local layer 6 cortical networks, we injected the Foxp2-Cre/TVA-G mice with libraries of ibRV at E16.5 using *in utero* injection and induced circuit labeling with TMP at E16.5, P0 and P7 for one week as described in Fig. 3. Since ibRV barcodes are random, they cannot be detected with probe-based spatial transcriptomics methods such as Merscope. Therefore, we utilized Open-ST, an unbiased spatial transcriptomics method with subcellular resolution^51^. Open-ST repurposes Illumina flow cells to generate high-resolution (0.6um) oligo capture surface for thin tissue sections. We generated Open-ST capture areas by depositing and sequencing a library of spatial barcodes (**Fig. 5A**) and utilized them to capture RNA from 10um brain tissue sections of P7 ibRV-infected brain. We generated two libraries to capture mRNA and rabies barcodes. We performed the analysis of the ibRV-Open-ST data by first segmenting the DAPI staining image and assigning spatially resolved transcripts to the segmented cells. Segmentation revealed that cell outlines matched the RNA-dense clusters both at the level of the entire tissue section and the level of individual cells, suggesting the single-cell accuracy of assignment of transcripts to spatial spots and cells (**Fig. 5B**). Unbiased clustering identified brain region and layer-enriched clusters, and neuronal subtype-specific marker genes demonstrated laminar and regional expression, including markers of cortical layer-specific excitatory neurons (ExNeu) (**Fig. 5C**). Additionally, expression of RV transcripts revealed widespread labeling with ibRV, demonstrating the efficiency of *in utero* labeling. Finally, we mapped the ibRV barcode sequences from the separate Open-ST library of the same tissue section onto the spatial coordinates shared with mRNA transcripts. We identified 3024 unique ibRV barcodes, with an average of 9 synaptic partners per barcodes captured in a single section. We reconstructed the cell type-specific single-cell wiring diagrams (**Fig. 5D**), proving the ability of the combination of ibRV and Open-ST to reveal the organization of specific brain circuits with single-cell and spatial resolution.

**Figure 5.**
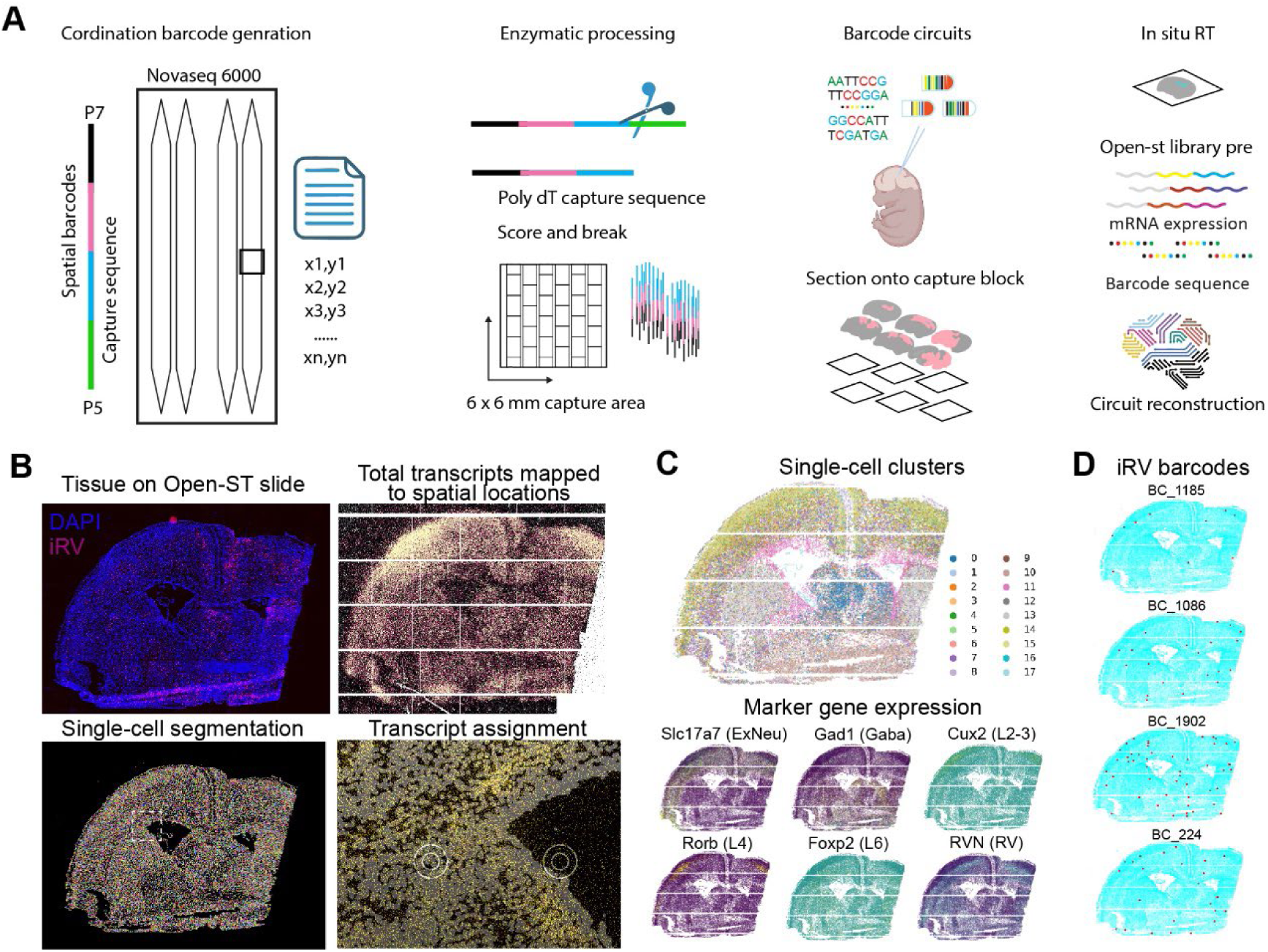
*In situ* single-cell sequencing combined with ibRV enables connectome analysis. (A) Repurposed flowcell used for *in situ* single-cell sequencing of ibRV-labeled brains. (B) Representative P7 cortical section showing fluorescence staining, single-cell segmentation, and transcript assignment. (C) Single-cell clustering with key cell-type markers. (D) Spatial distribution of example barcodes within the section.

## Discussion

scRNA-seq fundamentally transformed transcriptomics by overcoming the averaging effect of bulk sequencing, enabling the identification of rare cell states and molecular heterogeneity with unprecedented resolution^52–56^. Recent advances integrating spatially resolved methods such as *in situ* sequencing have further enhanced this power by embedding transcriptional information into tissue topology, revealing the cellular microenvironments and intercellular interactions underlying tissue function^51,57–60^. The brain, however, remains uniquely challenging due to its diverse cellular composition, dynamic developmental processes, and dense synaptic architecture. snRNA-seq has been particularly impactful in neuroscience because it preserves transcriptional states from complex neural tissue and enables access to deeply embedded or postmortem samples^61,62^. This approach has uncovered key mechanisms of neuronal differentiation, migration, and circuit maturation and revealed vulnerable cell populations associated with neurodevelopmental, psychiatric, and neurodegenerative disorders^61–68^. Yet, while molecular identity is crucial, it represents only one dimension of brain function: neuronal computation emerges from precise patterns of synaptic connectivity, which remain invisible to transcriptomic sequencing alone.

The convergence of rabies virus based synaptic tracing and high-throughput sequencing addresses this fundamental limitation^17–19^. Barcoded glycoprotein-deleted rabies virus, engineered to transfer RNA barcodes monosynaptically from starter neurons, allows direct measurement of wiring specificity alongside cell-state profiling. This emerging paradigm bridges molecular diversity and circuit architecture at single-cell resolution, enabling the reconstruction of synaptic inputs within heterogeneous neural populations. However, the practical impact of barcoded rabies systems has been constrained by one major biological bottleneck: cytotoxicity^21–29^. Even with optimized helper systems, rapid neuronal stress and degeneration can lead to premature loss of starter cells and incomplete mapping of their presynaptic partners, particularly in longer developmental studies.

Focusing on this limitation, we developed ibRV designed to reduce viral toxicity while preserving efficient monosynaptic transmission. By placing core viral replication factors, P and L^32^, under a molecular switch, we enabled temporal control of viral gene expression and observed markedly improved neuronal survival both *in vitro* and *in vivo*. This reduction in toxicity extended the feasible experimental window, permitting large-scale developmental tracing efforts that were previously unattainable. We leveraged this enhanced viability to perform embryonic (E16.5) labeling and combined ibRV readouts with *in situ* single-cell sequencing, capturing spatially resolved connectivity and transcriptional states across developing cortical circuits. Together, these innovations provide a powerful framework to interrogate how circuit motifs and cellular identities co-emerge during brain development.

Despite these accomplishments, we recognize that the results presented here reflect an early stage of technological deployment. Future optimization will focus on minimizing multiple infections per cell, increasing barcode diversity retention, and integrating computational strategies to refine circuit reconstruction accuracy. Additionally, extending ibRV to postnatal development and disease models, such as Alzheimer’s disease and autism spectrum disorder, will allow systematic investigation of how genetic risk and pathological processes reshape synaptic architecture over time.

## Methods

### Plasmid construction

The inducible rabies constructs were generated in the pSADΔG-mCherry backbone by inserting destabilizing domains at the N- or C-termini of the rabies N, P, and L open reading frames. An E. coli dihydrofolate reductase (ecDHFR)–based destabilizing domain was PCR-amplified from pBMN YFP-DHFR (Addgene, #29326). In parallel, a Shield-1–responsive FKBP12-derived DD cassette (sequence designed to be stabilized by Shield-1) was synthesized *de novo* (Integrated DNA Technologies, IDT). DD modules and junctions were assembled into pSADΔG-mCherry using the NEB HiFi DNA Assembly Master Mix (NEB, E5520) following the manufacturer’s instructions. In total, 24 constructs were generated: 12 with DHFR and 12 with DD, each fused to either the N- or C-terminal of N, P, or L proteins. Construct names reflect both the fusion position and protein target. For example, a construct with DHFR fused to the N-terminus of the P protein was named PnDHFR, while a construct with the DD fused to the N-termini of both P and L proteins was designated PLnDD. For pSADΔG-PnDHFR-H2B-mCherry, the H2B-mCherry sequence was PCR-amplified from pLenti6-H2B-mCherry (Addgene, #89766) and inserted into the pSADΔG-PnDHFR-mCherry backbone via HiFi DNA Assembly. All plasmids were verified by whole-plasmid Sanger sequencing (Eurofins Genomics). To construct a barcoded rabies virus plasmid library, template plasmids were linearized by targeting mCherry downstream region using two primers each containing a 10-nucleotide randomized barcode flanked by AscI or Pac1 restriction sites. Following PCR, products were digested with the corresponding restriction enzymes and circularized by ligation as previously described^18^. The ligation products were then transformed into electrocompetent *E. coli*, followed by outgrowth, large-scale colony expansion, and endotoxin-free plasmid purification to yield the final barcoded plasmid library. For barcode diversity assessment, 5–25 ng of the plasmid pool was combined with 25 μL of Q5® Hot Start High-Fidelity 2X Master Mix (NEB, M0494S), 2 μL of a UMI-containing oligonucleotide (see Table 1), and nuclease-free water to a final volume of 50 μL. The reaction was cycled under the following conditions: 98 °C for 3 min, 68 °C for 30 s, and 72 °C for 20 s. UMI-labeled products were selectively amplified (15 cycles) using primers incorporating Illumina P5 and P7 index adapters. The final amplicon libraries were sequenced on an Illumina NovaSeq X Plus platform using paired-end 150-cycle TruSeq chemistry. Two independent 10-bp barcode sequences were computationally extracted from each barcode cassette. To correct for PCR and sequencing errors, barcodes differing by a Hamming distance of 1 were collapsed during downstream analysis.

### Cell culture and viral transduction

HEK293T cells were obtained from the Duke Cell Culture Facility (CCF) and originally derived from the American Type Culture Collection (ATCC). Rabies virus production cell lines, including B7GG, BHK-EnvA, and HEK-TVA, were generously provided by Dr. Edward M. Callaway. Unless otherwise specified, all cell lines were cultured in Dulbecco’s Modified Eagle Medium (DMEM) supplemented with 10% fetal bovine serum (FBS) and 1% penicillin-streptomycin (P/S), and maintained at 37 °C in a humidified incubator with 5% CO₂. Routine mycoplasma testing was performed using the MycoStrip detection kit (InvivoGen, rep-mysnc-100). To generate the 293T7GG cell line for rabies virus rescue, HEK293T cells were transduced with lentiviral vectors encoding T7 RNA polymerase and EGFP-F2A-B19G. Following puromycin selection, GFP-positive cells were enriched by fluorescence-activated cell sorting (FACS) to establish a stable producer line.

### Virus production and purification

Rabies virus was produced in-house by transient transfection of B7GG cells with the rabies genome plasmid and a set of helper plasmids encoding B19 nucleoprotein (N), phosphoprotein (P), large polymerase (L), and glycoprotein (G) (pcDNA-SADB19N/P/L/G). Transfections were carried out using Lipofectamine™ 3000 and P3000 reagent (Thermo Fisher Scientific) in Opti-MEM for 8 hours, after which the medium was replaced with complete growth medium (DMEM + 10% FBS + 1% P/S). The following day, cells were transferred to an incubator set at 35 °C with 3% CO₂ to support viral rescue. When widespread red fluorescence was observed—indicating mCherry expression from the viral genome—the supernatant was harvested, filtered through a 0.45 μm membrane to remove debris, and titrated on 293T cells. Viral supernatants were aliquoted and stored at −80 °C. The inducible barcoded rabies virus (ibRV) library was generated similarly by transient transfection of 293T7GG cells with the barcoded rabies genome plasmid and helper plasmids (pcDNA-SADB19N/P/L/G; Addgene plasmids #100801, #100808, #100812, #100811). One day prior to transfection, 293T7GG cells were seeded at 30–40% confluency on si× 10-cm dishes coated with poly-D-lysine. Transfections were performed in Opti-MEM using Lipofectamine™ 3000 and P3000 for 8 hours, followed by medium replacement with complete DMEM. Cells were then moved to a 35 °C, 3% CO₂ incubator. Supernatants were collected once the majority of cells displayed red fluorescence, filtered (0.45 μm), and titrated on 293T cells. For pseudotyping, the filtered, crude CRV supernatant was used to infect BHK-EnvA cells at a multiplicity of infection (MOI) of 0.3. After 8 hours, infected cells were trypsinized, washed with PBS, and replated in fresh dishes. To remove residual unpseudotyped virus, cells were washed again with complete medium prior to further culture. EnvA-pseudotyped CRV was harvested from the supernatant 3–5 days post-infection. Viral titer and pseudotyping efficiency were assessed by infecting 293TVA and 293T cells plated in 24-well plates. For final preparation, pseudotyped viral supernatants underwent benzonase treatment to degrade residual contaminating nucleic acids, followed by ultracentrifugation and an additional spin through a 20% sucrose cushion. The viral pellets were resuspended in PBS (pH 7.4), aliquoted, titrated again, and stored at −80 °C until use.

### Barcode rabies virus library quality control

Genomic DNA from CRV-infected cells was used as template for targeted amplification of the sgRNA protospacer region. Primers (Table 1) flanking the protospacer sequence were used for PCR with Q5 Hot Start High-Fidelity DNA Polymerase (NEB, M0491L) for 10 cycles. PCR products were purified with AMPure XP beads (Beckman Coulter, A63881), and sequencing adapters were added by an additional 8–10 cycles of PCR with Q5 Hot Start to generate TruSeq Dual Index libraries. Libraries were pooled and sequenced on an Illumina NovaSeq 6000 platform with paired-end 150 bp reads (Novogene). Raw sequencing data were quality filtered using FastQC. Mutation profiles were then analyzed and visualized with CRISPResso2.

### Western blot analysis

Cells were washed with cold PBS and counted before lysed in Cell Lysis Buffer II (with protease inhibitors) on ice for 5 minutes. Total protein concentration was measured using the Bradford assay. SDS 6×loading buffer was added to the lysates, followed by boiling at 100°C for 10 minutes. Proteins were separated by SDS–PAGE (Bio-Rad, 4561036) and transferred to nitrocellulose membranes (Bio-Rad, 1620112). Membranes were blocked with 5% BSA in TBS-T (TBS with 0.1% Tween-20) for 30 minutes at room temperature, then incubated with primary antibodies (Table 2) at 4°C overnight. After three washes with TBS-T, membranes were incubated with HRP-conjugated secondary antibodies (Table 2) at room temperature for 1 hour. Protein bands were detected using the Clarity Max ECL Substrate (Bio-Rad, 1705062S) on a Bio-Rad imaging system.

### Transgenic mouse breeding and maintenance

B6.Cg-Gt(ROSA)26Sor^tm6(CAG-ZsGreen1)Hze^/J (Ai6, RRID:IMSR_JAX:007906) mice were breed homozygous and crossed with RphiGT for stereotaxic surgery. Male B6;129P2-Gt(ROSA)26Sor^tm1(CAG-RABVgp4,-TVA)Arenk^/J (RphiGT, RRID:IMSR_JAX:024708) were crossed with female CD-1® IGS mouse to improve maternal care and pup survival. Then male B6.Cg-Foxp2^tm1.1(cre)Rpa^/J mice (Foxp2-Cre, RRID:IMSR_JAX:030541), were crossed with female carrying the RphiGT allele to generate experimental animals hemizygous for both alleles. Offspring embryos or neonates were used for free hand (P0) or intro uterus cerebellum ventricular injection (E16.5). All mice were housed in the Duke Division of Laboratory Animal Resources (DLAR) barrier facility under a 12 h light/dark cycle, with ad libitum access to food and water. Breeding pairs were maintained on LabDiet® 5058, whereas non-breeding animals were fed LabDiet® 5053. Mice that underwent surgical procedures were provided with DietGel® 76A (ClearH2O, 72-07-5022) as a nutritional supplement to support recovery.

### Animal preparation and viral injection

All animal procedures complied with the NIH Guidelines for the Care and Use of Laboratory Animals and were approved by the Duke Institutional Animal Care and Use Committee (IACUC). For embryonic injections, timed pregnancies were established by housing one male with two females per cage. The following morning, females were checked for the presence of a vaginal plug, which was designated as embryonic day 0.5 (E0.5). Plug-positive females were then separated from the male, individually housed, and labeled for subsequent tracking of embryonic development. At the anticipated E10.5 stage, maternal body weight was measured to confirm pregnancy (>1.5g increase); females that did not gain weight were re-paired for breeding. On the day of surgery (E16.5), body weight was measured again to reconfirm pregnancy status. Pregnant females in good health were anesthetized with isoflurane and placed on a heating pad. The abdominal area was shaved and sterilized. Surgical procedures were performed as previously described^69,70^. Embryos were exposed, and the lateral ventricle of the cerebellum was targeted using anatomical landmarks (eye and lambda) to guide injection. A glass micropipette loaded with 1 μl virus premixed with Fast Green dye was inserted into the ventricle, and the solution was injected slowly. The pipette was held in place for 10 seconds post-injection to ensure complete delivery. Successful injection was confirmed by the presence and distribution of Fast Green within the ventricle. The number of embryos injected and non-injected littermates were recorded. After the procedure, the dam was returned to a clean cage with DietGel® 76A (ClearH2O, 72-07-5022) as nutritional support. Once the animal regained spontaneous movement, it was removed from the heating pad and returned to the housing rack. Dams were monitored daily for signs of distress and for the birth of pups. For postnatal day 0 (P0) injections, neonatal mice were anesthetized via hypothermia by placing them on ice overlaid with a latex glove until they no longer responded to toe pinch. The injection site was sterilized, and the target injection coordinate was determined as the point one-third of the distance between the eye and lambda, closer to lambda. Following 1 μl viral solution injection, pups were transferred to a heating pad for recovery and subsequently returned to their home cage once normal movement resumed.

For stereotaxic surgery, male and female mice (8–10 weeks old) were weighed before surgery and anesthetized with 3% isoflurane for induction. Animals were secured in a mouse adaptor on an Automated Stereotaxic Instrument (RWD, Model 71000) with a heating pad at 37 °C. Meloxicam (5 mg/kg) was administered subcutaneously for analgesia, the scalp was shaved and disinfected, and bupivacaine (0.25%, 2 mg/kg) was injected locally for additional analgesia. Throughout the procedure, anesthesia was maintained with 1.5% isoflurane delivered through a nose cone at 0.5 L/min. Injection sites were identified using coordinates from the Paxinos and Franklin Mouse Brain Atlas (2nd Edition). A burr hole (∼0.6 mm) was drilled, and viral vectors were delivered with pulled glass micropipettes using a micropump (WPI). The micropipette was left in place for 5 min after injection. The craniotomy was sealed with bone wax, and the scalp was sutured. Following recovery, mice were returned to clean cages. Post-surgical care included daily meloxicam injections on days 1 and 2. Mice exhibiting significant weight loss were euthanized according to approved protocols.

### Histological analysis

Mice were anesthetized with isoflurane and perfused transcardially with cold PBS followed by 4% paraformaldehyde (PFA). Brains were dissected, fixed in 4% PFA overnight at 4 °C, and transferred to 30% sucrose in PBS 1-2 days for cryoprotection. For neonatal mice younger than P6, euthanasia was performed via hypothermia, followed by immediate brain dissection and immersion fixation in 4% PFA overnight at 4 °C. The brains were then cryoprotected and processed as described above. Tissues were embedded in OCT & Sucrose (30%) compound (3:2 v/v) and sectioned on a cryostat. For free-floating immunostaining, sections were permeabilized with 0.4% Triton X-100 in PBS for 5 min and blocked with 3% BSA in PBS for 1 h at room temperature. Sections were incubated with primary antibodies at 4 °C overnight, washed three times with PBST (PBS + 0.1% Tween-20), and then incubated with secondary antibodies for 2 h at room temperature. After washing, sections were counterstained with DAPI and mounted. Images were acquired using a Zeiss LSM 880 confocal microscope.

### Flow Cytometry

For flow cytometric analysis, cells were dissociated using trypsin, washed with PBS, and passed through a 40 µm cell strainer. Flow cytometry was performed using a BD LSRFortessa™ Cell Analyzer to assess viral infection efficiency (mCherry expression) and editing efficiency (loss of EGFP fluorescence). To generate stable cell lines expressing fluorescent proteins, cells were transduced with lentiviral vectors and subsequently sorted using a Sony MA900 cell sorter equipped with a 100 µm microfluidic chip.

### *In situ* single-cell sequencing with repurposed flowcell

*In situ* single-cell RNA sequencing was performed using the Open-ST protocol^51^. Briefly, a NovaSeq 6000 flow cell was pre-sequenced by Novogene to obtain spatial coordinate information for subsequent indexing, following the methodology described in the original Open-ST publication. After sequencing, the flow cell was rinsed with ultrapure water, sealed, and shipped to our laboratory for downstream processing. Surface functionalization and array preparation were carried out according to the Open-ST protocol. The treated flow cell was sectioned into 6 × 6 mm capture areas, dried at −20 °C, and stored until use.

Fresh-frozen, OCT-embedded brains from neonates previously infected with barcoded rabies virus were cryosectioned at a thickness of 10 μm. Sections were screened under a fluorescence microscope for anatomical landmarks and H2B-mCherry expression. After placement on the flow cell capture area, tissue sections were DAPI-stained and imaged before RNA capture. RNA and rabies barcode transcripts were simultaneously captured using the Open-ST workflow. Following cDNA synthesis and library preparation, 10% of the total library input was used for targeted amplification of the barcode cassette to generate a dedicated barcode sub-library. Both the whole-transcriptome library and the barcode sub-library were sequenced on an Illumina NovaSeq X Plus platform using paired-end 150 bp reads (PE150), achieving a sequencing depth exceeding 600 million reads per section. Data processing and analysis followed the standard Open-ST computational pipeline, with an additional step to match individual cells to rabies barcodes based on the pre-acquired spatial coordinate information.

## Data and materials availability

All data supporting the findings of this study are included in the main text and supplementary materials. Key plasmids will be made available through Addgene.

## Code availability

All code is available at Github at https://github.com/velmeshevlab/CRV. The single-cell sequencing datasets will be deposited at SRA and GEO.

## Acknowledgments

We thank all members of the Dmitry Velemshev lab for helpful advice and discussions. We thank the Duke Core Facilities for assistance with animal, flow cytometry and imaging. We thank Dr. Z. Josh Huang and Baoxia Han for their guidance on transgenic mouse genotyping and breeding. This study is supported by NIH R00 grant R00MH121534, DP1 grant DP1DA063507, Duke University’s Whitehead Scholarship, as well as a generous donation by David Dolby to DV. ZZ is supported by the postdoctoral fellowship from the Ruth K. Broad Biomedical Research Foundation.

## Author contributions

Conceptualization: ZZ, BRD, DLS, DV

Methodology: ZZ, BRD, EM, DGS, ZJH, DLS, DV

Investigation: ZZ, BRD, LH, DA, SD, EM, ZY, NR, KSK, DS, ZJH, DV

Visualization: ZZ, DV

Funding acquisition: DV

Project administration: DV

Supervision: DV

Writing – original draft: ZZ, DV

Writing – review & editing:

## Competing interests

ZZ and DV are inventors of the inducible rabies virus technology listed in a provisional patent USSN 63/910,441

## Supplementary Materials

Figs. s1 to s6

Tables 1 and 2

**Figure s1.**
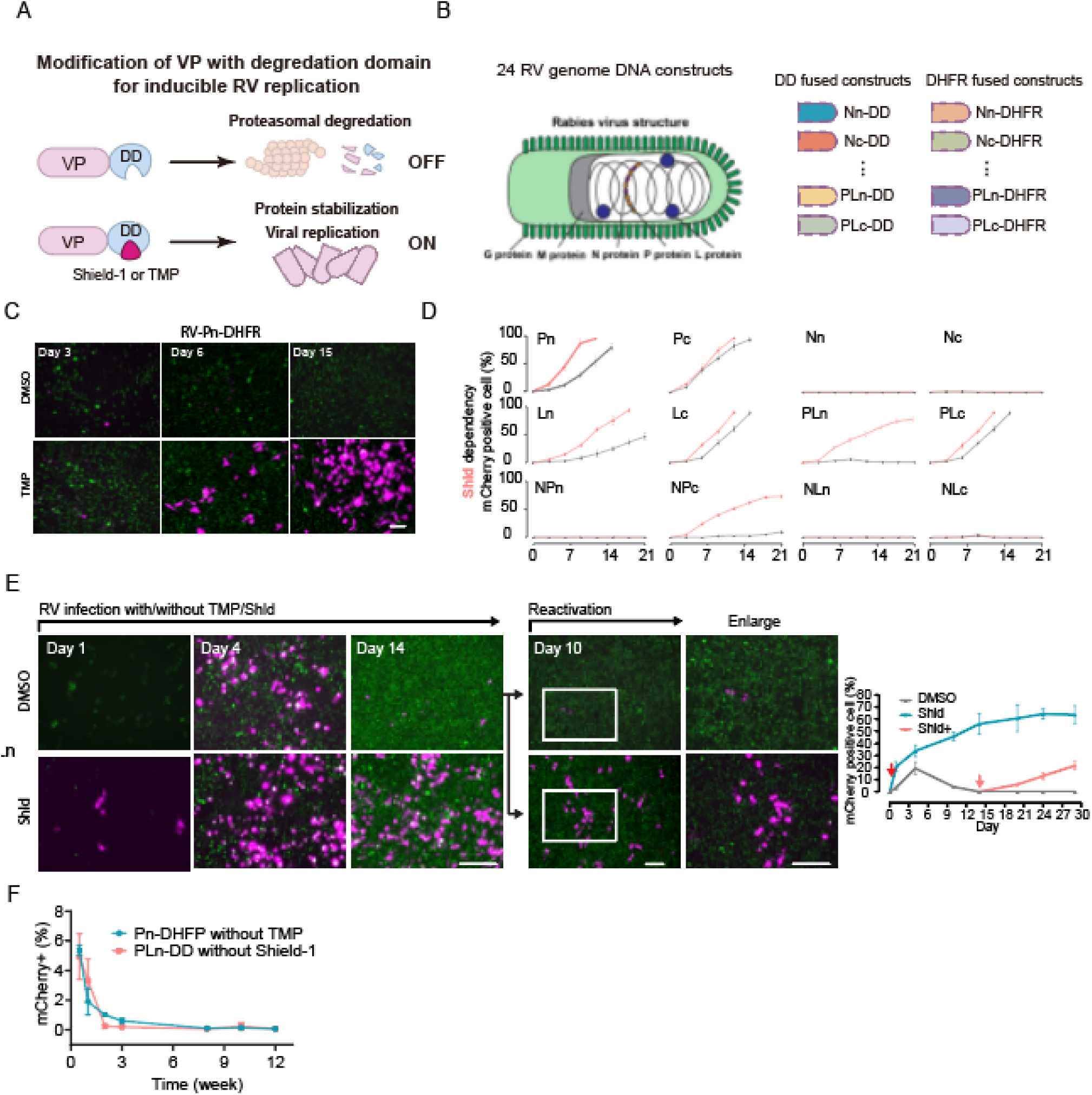
iRV through fusion with molecular switches achieve inducible proliferation.

**Figure s2.**
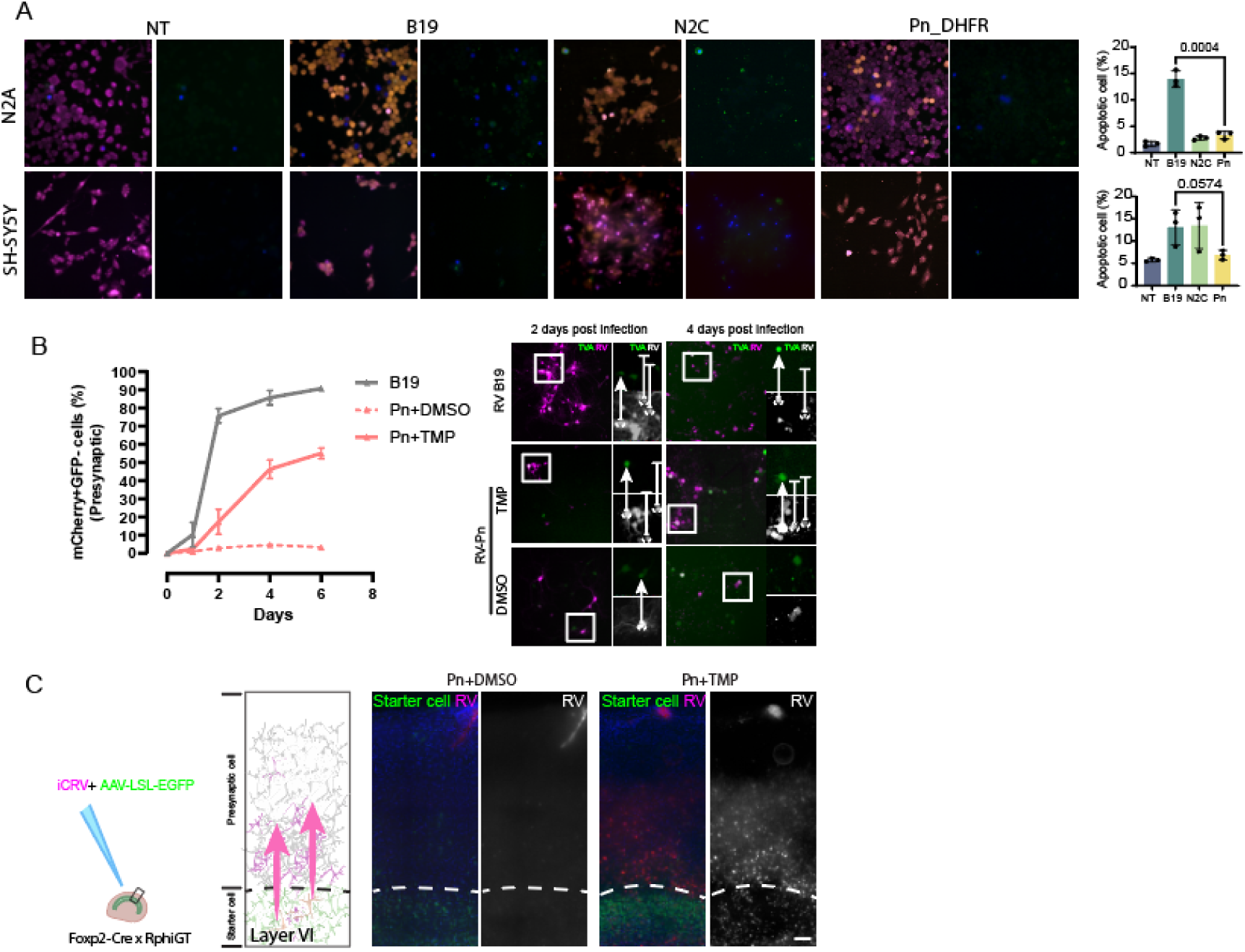
iRV retains transsynaptic labeling capability while exhibiting reduced toxicity.

**Figure s3.**
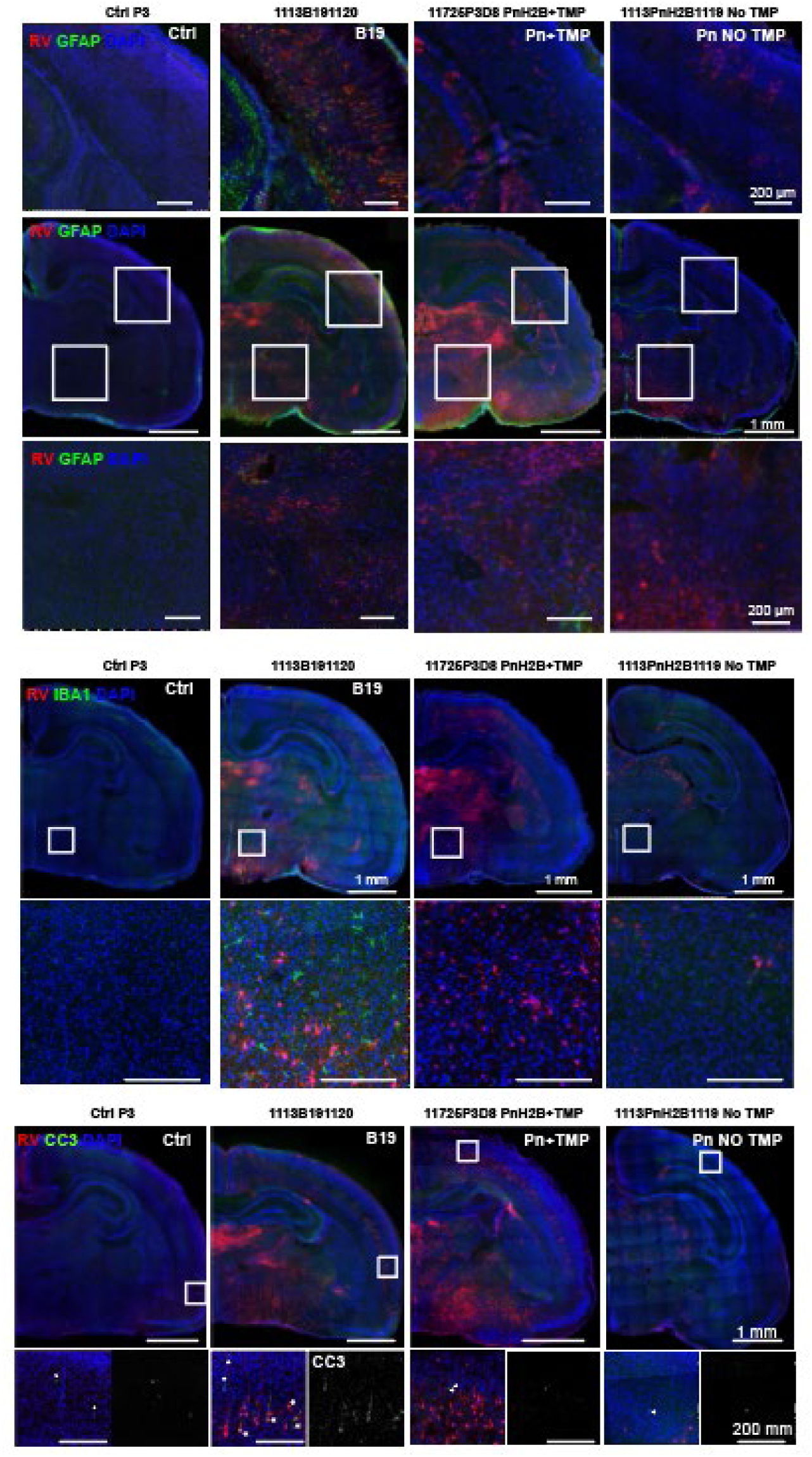
Intro-uterine injection of iRV eliciting lower levels of immune response and apoptosis than B19.

**Figure s4.**
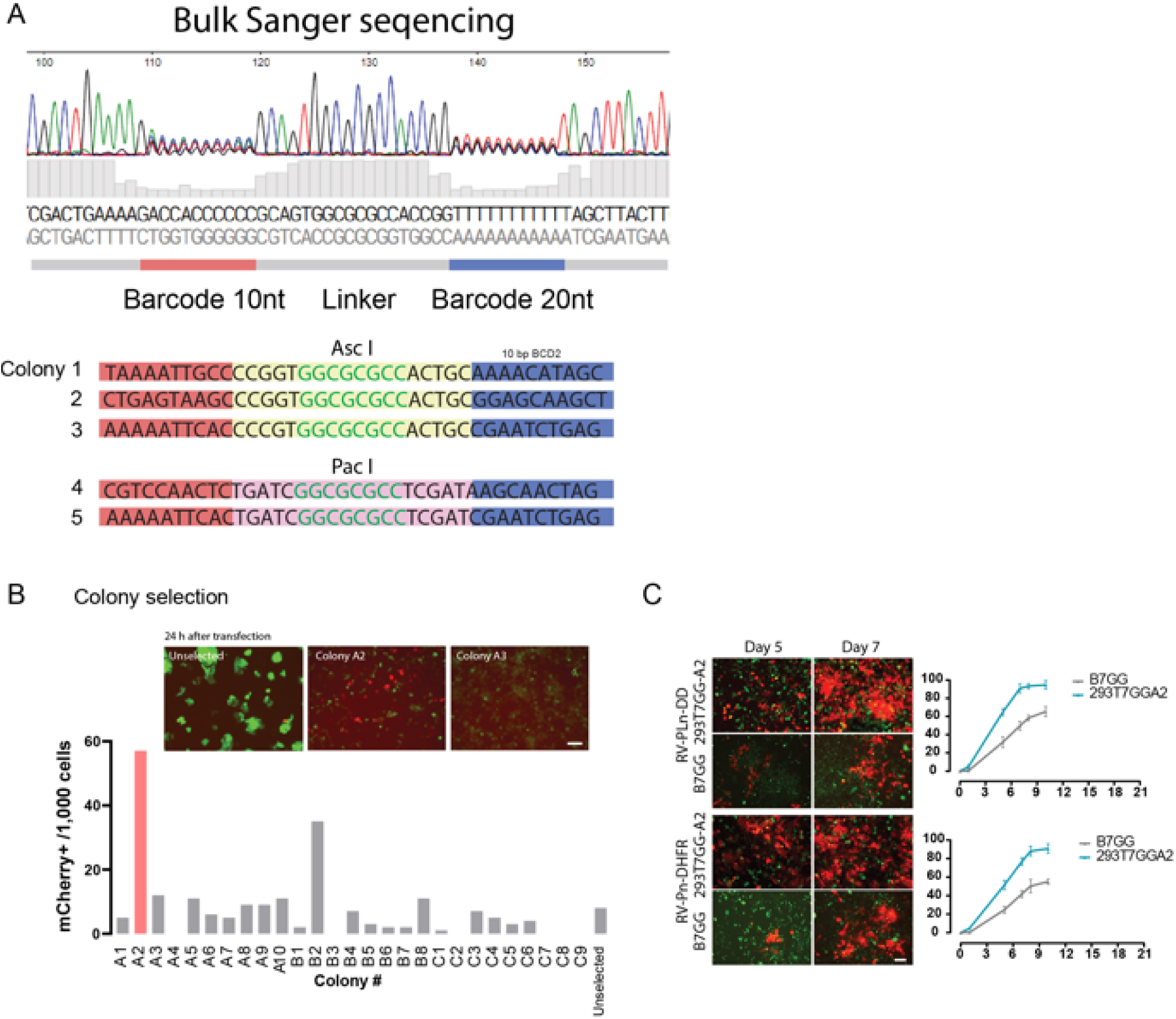
Optimization of ibRV production to improve yield and quality.

